# REM disruption and REM Vagal Activity Predict Extinction Recall in Trauma-Exposed Individuals

**DOI:** 10.1101/2023.09.28.560007

**Authors:** Cagri Yuksel, Lauren Watford, Monami Muranaka, Emma McCoy, Hannah Lax, Augustus Kram Mendelsohn, Katelyn I. Oliver, Carolina Daffre, Alexis Acosta, Abegail Vidrin, Uriel Martinez, Natasha Lasko, Scott Orr, Edward F. Pace-Schott

## Abstract

Accumulating evidence suggests that rapid eye movement sleep (REM) supports the consolidation of extinction memory. REM is disrupted in PTSD, and REM abnormalities after traumatic events increase the risk of developing PTSD. Therefore, it was hypothesized that abnormal REM in trauma-exposed individuals may pave the way for PTSD by interfering with the processing of extinction memory. In addition, PTSD patients display reduced vagal activity. Vagal activity contributes to the strengthening of memories, including fear extinction memory, and recent studies show that the role of vagus in memory processing extends to memory consolidation during sleep. Therefore, it is plausible that reduced vagal activity during sleep in trauma-exposed individuals may be an additional mechanism that impairs extinction memory consolidation. However, to date, the contribution of sleep vagal activity to the consolidation of extinction memory or any emotional memory has not been investigated. To test these hypotheses, we examined the association of extinction memory with REM characteristics and REM vagal activity (indexed as heart rate variability) in a large sample of trauma-exposed individuals (n=113). Consistent with our hypotheses, REM disruption was associated with poorer physiological and explicit extinction memory. Furthermore, higher vagal activity during REM was associated with better explicit extinction memory, and physiological extinction memory in males. These findings support the notion that abnormal REM may contribute to PTSD by impairing the consolidation of extinction memory and indicate the potential utility of interventions that target REM sleep characteristics and REM vagal activity in fear-related disorders.

## INTRODUCTION

PTSD is characterized by excessive fear responses to cues associated with a traumatic event. It is hypothesized that this is due to dysfunction in fear extinction, the process whereby fear response is gradually reduced after multiple exposures to a conditioned stimulus (CS) without reinforcement (Milad and Quirk 2012; Zuj et al. 2016). After a traumatic event, fear extinction is achieved by repeated exposure to reminders of the traumatic event without the feared outcome. This mechanism is suggested to be the basis of exposure therapy for PTSD and other anxiety disorders (Craske et al. 2014).

Fear extinction is not simply an erasure of fear but relies on new learning and memory. While the preponderance of studies does not show impaired extinction learning in PTSD (reviewed in (Bottary et al. 2023; Pace-Schott et al. 2023), accumulating evidence suggests that impaired retention of extinction memory, acquired after trauma exposure (Milad et al. 2008), may be a critical mechanism that leads to PTSD (Helpman et al. 2016; Milad et al. 2008; Milad et al. 2009; Milad and Quirk 2012; Shvil et al. 2014; Suarez-Jimenez et al. 2020; Wicking et al. 2016). Sleep supports the consolidation of different types of memories, which stabilizes and integrates them, and enhances their retrieval (Klinzing et al. 2019). Recent evidence suggests that this extends to extinction memory, with rapid eye movement sleep (REM) implicated as an important sleep stage involved in its consolidation (Davidson and Pace-Schott 2020). In healthy people, more REM has been shown to be associated with better extinction recall (Bottary et al. 2020; Menz et al. 2016; Pace-Schott et al. 2014; Spoormaker et al. 2010), and REM deprivation leads to impaired extinction memory (Spoormaker et al. 2012), suggesting the possibility of a causal role for this sleep stage. Several studies and meta-analyses find REM abnormalities in PTSD (Kobayashi et al. 2007; Richards et al. 2020; Zhang et al. 2019), which also have been shown to be associated with the risk of developing PTSD after trauma (Mellman et al. 2002; Mellman et al. 2004; Mellman et al. 2007). Therefore, it is postulated that sleep disruption, specifically of REM, may predispose traumatized individuals to develop PTSD symptoms by interfering with extinction memory (Colvonen et al. 2019; Pace-Schott et al. 2023). Supporting this hypothesis, in a sample overlapping that of the current report, we showed that prefrontal activations during extinction recall were associated with REM density during the night before extinction recall (Seo et al. 2022). However, much remains to be learned regarding the association between REM and extinction memory in trauma-exposed individuals. One preliminary study (Straus et al. 2018) did not find any association of REM with extinction recall in PTSD; however, the small sample size (n=13) may explain this negative finding. This study’s sample was also limited to those with PTSD diagnoses and, therefore, did not provide insight into this association in individuals with subthreshold PTSD symptoms. To address this gap in knowledge, we examine here the association of REM measures on the night following extinction learning with extinction recall the following day in a large sample of trauma-exposed individuals. We hypothesized that disruptions in REM sleep would be associated with impaired extinction recall (Hypothesis 1).

PTSD is characterized by abnormal autonomic nervous system function, including reduced parasympathetic nervous system (PNS) activity (Seligowski et al. 2022). PNS activity can be estimated by measuring heart rate variability (HRV), the variation in the interval between successive heartbeats. Specific HRV measures in the frequency (high frequency; HF-HRV) and time (root mean square of successive differences in the R–R interval; RMSSD) domains reflect control of heart rate by the vagus nerve, the main component of PNS outflow from the CNS (Laborde et al. 2017; Shaffer and Ginsberg 2017). Meta-analyses show that patients with PTSD display reduced vagal activity, as indexed by HF-HRV and RMSSD, during wakefulness (Chalmers et al. 2014; Ge et al. 2020; Schneider and Schwerdtfeger 2020), and in one study, lower baseline HF-HRV predicted the severity of later post-traumatic stress symptoms (Pyne et al. 2016). Similar to wakefulness, a small number of recent studies in sleep find reduced vagal activity in PTSD (Kobayashi et al. 2014), including in specific sleep stages (Daffre et al. 2023; Ulmer et al. 2018). Additionally, in one study, HF-HRV across actigraphically measured sleep was associated with PTSD symptom severity (Rissling et al. 2016).

Converging evidence indicates that vagal activity facilitates emotional processing and memory, likely via its afferents to the brainstem nuclei, which in turn modulate the activity of multiple neurotransmitter systems and relevant brain regions, including the prefrontal cortex (PFC), amygdala, and hippocampal formation (Broncel et al. 2020; McGaugh 2013). Consistent with this, high vagal tone is associated with better extinction learning in humans (Jenness et al. 2019; Pappens et al. 2014; Wendt et al. 2015), and enhancing vagal activity improves extinction learning and retention of extinction memory in both rodents (Noble et al. 2017; NobleMeruva et al. 2019; Pena et al. 2014; Pena et al. 2013; Souza et al. 2022) and humans (Burger et al. 2019; Burger et al. 2016; Szeska et al. 2020). In addition, a separate line of recent studies shows that vagal activity during sleep, as indexed by HF-HRV, independently contributes to memory consolidation over and above contributions by sleep architecture and sleep-related neural oscillations (Whitehurst et al. 2016; Whitehurst et al. 2022). However, to date, no study has examined the role of vagal activity in the sleep-dependent consolidation of extinction memory in any population. We thus hypothesized that lower HF-HRV in REM during the night following extinction learning would independently predict lower extinction recall in trauma-exposed individuals (Hypothesis 2). If this hypothesis is supported, it would further underscore vagal activity as a promising target for enhancing extinction memory through readily accessible interventions (Jung et al. 2024; Siepmann et al. 2022) in the aftermath of trauma exposure.

## METHODS

### Participants

A total of 139 participants, aged between 18 and 40, were recruited from the greater Boston metropolitan area using online and posted advertisements. All participants reported experiencing a DSM-5 criterion-A traumatic event (“index trauma”) in the past two years, except within the last month. Further inclusion and exclusion criteria can be found in previous publications (Daffre et al. 2023; Denis et al. 2021; Seo et al. 2022) and Supplementary Materials. Twenty-six participants were excluded from all analyses because of missing or unusable EEG or ECG data (final n=113). Psychiatric history was ascertained using the Structured Clinical Interview for DSM-IV-TR for Non-Patients (SCID-I/NP) (First et al. 2007). The Clinician-Administered PTSD Scale for DSM-5 (CAPS-5; (Weathers et al. 2018)) was used to assess post-traumatic stress symptom severity (see **Figure S1** for its distribution in the sample). This study followed a Research Domain Criteria (RDoC) (Insel et al. 2010) design in which dimensional rather than categorical measures were targeted, and PTSD diagnoses were established post-hoc from diagnostic evaluations rather than by assigning subjects to PTSD and non-PTSD groups. 49.6% (n=56) of participants met diagnostic criteria for PTSD. The Quick Inventory of Depressive Symptomatology, Self-Report (QIDS-SR) (Rush et al. 2003), and the Pittsburg Structured Clinical Interview for Sleep Disorders (SCID-SLD) (Stocker et al. 2017) were administered to evaluate depressive and sleep disorder symptoms, respectively. Demographic and clinical characteristics of the final sample are displayed in **Table S1**. All participants provided written consent to participate in the study and were paid for their participation. All procedures were approved by the Partners Healthcare Institutional Review Board.

### Procedures

Participants completed an approximately 2-week sleep assessment period during which they wore an actigraph (Actiwatch-2, Philips Respironics) and filled out a daily sleep and nightmare diary. Approximately midway through the 14-day period, they underwent a combined sleep-disorders-screening and acclimation night of ambulatory polysomnography (PSG) recording. This was followed by a “baseline” overnight PSG recording. Starting in the evening immediately after the baseline night, participants completed a 2-day fear conditioning and extinction protocol with simultaneous fMRI recordings (Milad et al. 2013). Between fMRI sessions, participants completed a third (“consolidation”) night of ambulatory PSG. Details of the PSG methods are included in the Supplementary Methods.

### Fear conditioning, extinction learning, and extinction recall

A validated 2-day paradigm (Milad et al. 2013; Milad et al. 2007) was used to probe fear conditioning and extinction during ongoing fMRI recording. This protocol consisted of 4 phases, with Habituation, Fear Conditioning, and Extinction Learning phases taking place on the first day and Extinction Recall 24 hours later. Images of a colored desk lamp (red, yellow, or blue) appearing in a conditioning and extinction-context background served as conditioned stimuli (CS). During fear conditioning, a mild electric shock was paired with the presentation of 2 differently colored lamps (CS+) but not a third color (CS-). During extinction learning, one CS+ (CS+E) but not the other (CS+U) was extinguished using presentations without any electric shock. During each phase, physiologic reactivity for each trial was indexed using skin conductance response (SCR), a measure of sympathetic activity (Dawson et al. 2007). Negative SCRs were coded as zero (Lonsdorf et al. 2017), and then all values were square root transformed. “Non-conditioners” were defined as those who exhibited less than two non-square-root transformed SCR responses to either of the two CS+s that were equal to or exceeding .05 µS during the Fear Conditioning phase (Orr et al. 2000). Thirty-one non-conditioners were further excluded from SCR analyses based on these criteria. In addition, SCR data for 3 participants were lost. Differential SCR (SCRd) was calculated by subtracting from the SCR to each CS+ the SCR from its closest temporally corresponding CS-to adjust for nonspecific reactivity to the nonreinforced CS- (Bottary et al. 2020; Lonsdorf et al. 2019).

Immediately following each phase except Habituation, participants verbally reported shock expectancy for the first and last presentations of each CS (i.e., colored light) appearing in that phase on a scale from 1 (“not expecting a shock at all”) to 5 (“expecting a shock very much”). Expectancy ratings for 7 participants were not included in the analyses, because they did not complete the experiment or because of missing data. Further details of the protocol and skin conductance monitoring are provided in previous publications (Bottary et al. 2020; Pace-Schott et al. 2009) and in Supplementary Materials.

### Extinction Recall Variables

To examine the association of physiologically expressed and subjective extinction recall with sleep and HRV measures, we used a Differential Extinction Retention Index (dERI) (Lonsdorf et al. 2019) and a Subjective Extinction Retention Index (sERI) (Bottary et al. 2020), respectively. dERI was calculated as: [(Average of the SCRds of the first 4 CS+E presentations at Extinction Recall phase/maximum SCRd to the “to-be” CS+E during the Fear Conditioning phase) x 100] (Lonsdorf et al. 2019). Only the first 4 CS+E trials from the Extinction Recall phase were included in this calculation in order to avoid confounding recalled extinction with new extinction learning. Higher dERI reflected lower extinction memory. sERI was calculated as: Expectancy to the first CS+E in Extinction Recall/mean expectancy for the last of each CS+ during Fear Conditioning × 100. Larger sERI indicated lower extinction memory.

### Sleep Measures

The PSG data analyzed in the present study were recorded during the consolidation night. For regression analyses, we selected the REM measures that have been associated with PTSD, including %REM (Zhang et al. 2019), REM density (REMD) (Kobayashi et al. 2007), REM latency (REML) (Mellman et al. 2014), and REM fragmentation (REMF) (Breslau et al. 2004; Habukawa et al. 2018; Insana et al. 2012; Lipinska and Thomas 2019; Mellman et al. 2002; Saguin et al. 2021). Percent time spent in each sleep stage (%N1-N3, %REM) was computed as a percentage of total sleep time (TST). REMD is the number of rapid eye movements per minute of REM sleep and was calculated using an automatic algorithm (Yetton et al. 2016). REML was calculated as the number of minutes occurring after sleep onset before the first REM epoch. REMF was calculated as the average duration of REM segments (Mellman et al. 2002). REM segments were defined as continuous REM from the start of at least 1 minute of REM to the onset of at least 1 minute of non-REM or wake (Mellman et al. 2002).

### Heart Rate Variability

HRV was calculated in continuous ECG segments of ≥5 minutes within REM using Kubios HRV premium software (Kubios Oy, Kuopio, Finland). Prior to analysis, ECG traces were visually inspected for physiological and technical artifacts (Laborde et al. 2017). Misplaced or ectopic beats were manually corrected if possible. Very low to low, or occasionally medium, filter was applied, if necessary, after correction (https://www.kubios.com/hrv-preprocessing/). Segments that did not provide reliable estimates due to excessive artifacts were removed. Twenty-three participants were excluded because of artifacts, not having ≥5 minute REM segments, or because their records were missing. High frequency (HF; .15-.4 Hz) absolute power (HF[ms^2^]) was used as a predictor variable in the primary analyses because it was shown to be associated with sleep-dependent memory processing (Whitehurst et al. 2016). We also carried out the same analyses with RMSSD, another measure of vagal activity (Shaffer and Ginsberg 2017). For frequency domain analyses, autoregressive method was used, with model order set at 16 (Laborde et al. 2017). Data was transformed by their natural logarithm (Laborde et al. 2017).

### Statistical Analyses

Associations were explored between extinction recall indices (dERI and sERI), and age, sex, months since index trauma, scores of symptom scales (CAPS-5 and QIDS), and medication use as a binary variable. For correlations, Pearson and Spearman’s correlations were used depending on the distribution of variables. Student’s t-tests were used for comparisons between sexes and between medication users and non-users.

We used linear regression models to test whether REM sleep variables predicted extinction recall (dERI and sERI; Hypothesis 1). The models included %REM, REMD, REML, and REMF as predictor variables. Sex and medication use were also included as predictors as these factors have been shown to modulate fear extinction and its retention (Bouton et al. 1990; Gunduz-Cinar et al. 2016; Milad et al. 2006).

Next, hierarchical regression analyses were used to test the hypothesis that vagal activity would improve the predictions of dERI and sERI above and beyond REM sleep measures alone (Hypothesis 2). For this purpose, two models were built. Model 1 included sex, medication use and REM measures. HF[ms^2^] was added as an additional predictor in Model 2. Because missing HF[ms^2^] data further limited the number of participants that could be included in the analysis, only REM variables that were significant predictors in the regression analysis that tested Hypothesis 1 were included. Results for the models that included all REM variables and RMSSD instead of HF[ms2] were also reported for completeness.

In all regression analyses, partial regression plots and a plot of studentized residuals against the predicted values indicated that assumptions of linear relationship were met. Variance inflation factors were within acceptable ranges (1.1-1.4), indicating that there was no confounding multicollinearity. For dERI, the distribution histogram and P-P plot of standardized residuals showed a non-normal distribution. In addition, plotting standardized residuals and predictors revealed heteroscedasticity. Therefore, dERI was log-transformed. After transformation, normal distributions of residuals and homoscedasticity were confirmed.

For all variables, outliers ± 3 standard deviations from the mean were removed. All analyses were two-tailed. The significance level for hypothesis testing (α) was set at .05.

## RESULTS

### Association of Extinction Recall Indexes with Demographic and Clinical Measures

Age was inversely correlated with sERI (r_s_= -.27, p=.006), while it was not significantly correlated with dERI (r_s_= .12, p=.305). There were no differences between sexes in dERI or sERI (t(76)=.233, p=.817 and t(104)=.116, p=.250, respectively). There were no significant correlations between either extinction recall index and the number of months since index trauma (dERI: r_s_= -.03, p=.796; sERI: r_s_= -.04, p=.659), or CAPS-5 (dERI: r_s_= .11, p=.333; sERI: r_s_=- .02, p=.831) or QIDS (dERI: r_s_= .13, p=.350; sERI: r_s_= -.04, p=.692) scores.

### Regression Analyses Predicting dERI and sERI

Measures of REM characteristics and sleep architecture are displayed in **Table 1**. The multiple regression model with dERI as the dependent variable included sex, medication use, %REM, REMD, REML, and REMF as predictor variables (N=67). The overall model explained a significant proportion of the variance (Adj. R^2^=.14, p=.021), with sex, **%**REM, REMD, and REML emerging as significant predictors (**Table 2; Figure S2**). Poorer physiological extinction recall was associated with the male sex (B[SE]=-.71[.34], p=.041), less time in REM sleep (%REM; B[SE]=-.06[.03], p=.018), increased REMD (B[SE]=.14[.05], p=.009), and shorter REML (B[SE]=-.01[.003], p=.002).

**Table 1.** Sleep and HRV measures in the participants. Data is displayed as mean ± standard deviation (minimum - maximum). HF[ms^2^]: Absolute power of high frequency heart rate variability; %N1-REM: Proportion of the sleep stage to the total sleep time; REMD: REM density; REMF: REM fragmentation; REML: REM latency; RMSSD: Root mean square of successive differences.

|  |  |
| --- | --- |
| **REMD** | 5.1±3.1 (.62-13.0) |
| **REML** | 102.3±45.9 (0-238.0) |
| **REMF** | 9.8±4.0 (2.8-23.4) |
| **%N1** | 5.7±3.8 (0.8-27.0) |
| **%N2** | 54.5±9.3 (26-81.7) |
| **%N3** | 21.7±10.6 (1.7-63.3) |
| **%REM** | 18.3±6.5 (2.0-35.0) |
| **HF[ms<sup>2</sup>]** | 1198.0±1069.5 (39.2-4788.6) |
| **RMSSD** | 54.3±26.8 (13.0-128.3) |

**Table 2.** Linear regression analysis for physiological extinction recall (dERI; N=67). Sex, %REM, REMD and REML were significant predictors. Note that smaller dERI denotes better extinction recall. %REM: Proportion of REM sleep to the total sleep time; REMD: REM density; REMF: REM fragmentation; REML: REM latency.

| ANOVA |  |  |  |  |  |  |  |  |  |  |
| --- | --- | --- | --- | --- | --- | --- | --- | --- | --- | --- |
| Predictors | B | SE | $\beta$ | t | p | 95% CI | | Adj. R<sup>2</sup> | F | p |
| Sex | -.714 | .341 | -.279 | -2.092 | .041 | -1.397 | -.031 | .135 | 2.716 | .021 |
| Medications | .113 | .372 | .038 | .304 | .762 | -.630 | .856 |  |  |  |
| %REM | -.061 | .025 | -.326 | -2.427 | .018 | -.111 | -.011 |  |  |  |
| REMD | .141 | .052 | .365 | 2.701 | .009 | .036 | .245 |  |  |  |
| REML | -.011 | .003 | -.421 | -3.167 | .002 | -.018 | -.004 |  |  |  |
| REMF | .036 | .031 | .153 | 1.155 | .253 | -.026 | .098 |  |  |  |

A similar multiple regression model for sERI was carried out (N=94). In our sample, age was associated with sERI (see Results), therefore it was also included in the model. This model also explained a significant proportion of the variance (Adj. R^2^=.09, p=.031). In this model, REMD was the only significant predictor (**Table 3; Figure S3**), and as with dERI, increased REMD was associated with poorer subjective extinction recall (B[SE]=2.80[.94], p=.004).

**Table 3.** Linear regression analysis for subjective extinction recall (sERI; N=94). REMD was the only significant predictor. Note that smaller sERI denotes better extinction recall. %REM: Proportion of REM sleep to the total sleep time; REMD: REM density; REMF: REM fragmentation; REML: REM latency.

|  |  |  |  |  |  |  |  | ANOVA |  |  |
| --- | --- | --- | --- | --- | --- | --- | --- | --- | --- | --- |
| Predictors | B | SE | $\beta$ | t | p | 95% CI | | Adj. R<sup>2</sup> | F | p |
| Age | -.512 | .629 | -.084 | -.814 | .418 | -1.763 | .739 | .092 | 2.345 | .031 |
| Sex | -2.903 | 6.842 | -.046 | -.424 | .672 | -16.503 | 10.698 |  |  |  |
| Medications | -12.574 | 7.632 | -.175 | -1.648 | .103 | -27.747 | 2.598 |  |  |  |
| %REM | .238 | .527 | .052 | .452 | .653 | -.810 | 1.286 |  |  |  |
| REMD | 2.801 | .936 | .331 | 2.994 | .004 | .942 | 4.661 |  |  |  |
| REML | .087 | .072 | .136 | 1.209 | .230 | -.056 | .229 |  |  |  |
| REMF | .528 | .653 | .090 | .808 | .421 | -.771 | 1.827 |  |  |  |

### Hierarchical regressions with HF HRV predicting dERI and sERI

Average HRV values are displayed in **Table 1**. For the hierarchical regression analyses with dERI as the dependent variable (N=61), sex, medication use, and REM sleep measures (%REM, REMD, and REML) were included in the first model. HF[ms2] was included as an additional predictor in the second model. Visual inspection of the data suggested that the association of HF[ms^2^] with dERI might be different between sexes. Therefore a sex × HF[ms^2^] interaction was also added to the second model. The final model was significant (Adj. R^2^=.17, p=.018; **Table 4**). There was a trend for an increase in the variance accounted for by the second model (ΔR^2^=.08, p=.074). HF[ms^2^] and the sex × HF[ms^2^] interaction were significant predictors (B[SE]=- .42[.20], p=.042 and B[SE]=.65[.29], p=.030, respectively. We therefore probed this interaction by examining the simple slopes, which showed that the effect of HF[ms^2^] on dERI was significant for males (B[SE]=-.42[.20], p=.042) but not for females (B[SE]=.23[.21], p=.263), and the effects were different between sexes (p=.031).

**Table 4.**
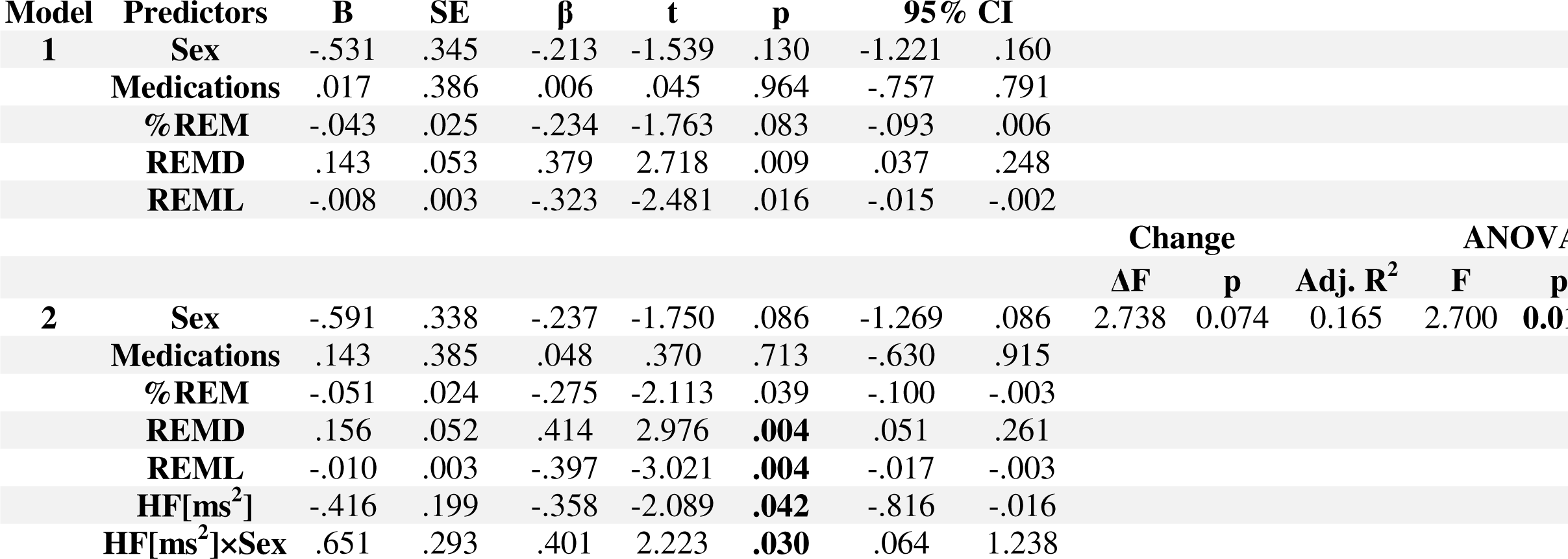
Hierarchical regression analysis for physiological extinction recall (dERI; N=61). HF[ms^2^] and HF[ms^2^]×Sex interaction were significant predictors. Note that smaller dERI denotes better extinction recall. HF[ms^2^]: Absolute power of high frequency heart rate variability; %REM: Proportion of REM sleep to the total sleep time; REMD: REM density; REML: REM latency.

A similar hierarchical regression analysis was carried out for sERI (N=85). The first model included age, sex, medication use, and REMD. HF[ms^2^] was added to the predictors in the second model. Addition of HF[ms^2^] led to a significant increase in the variance accounted for by the model (ΔR^2^=.06, p=.018), indicating that vagal activity contributed to the prediction of subjective extinction recall above and beyond REM sleep measures. The final model was significant (Adj. R^2^=.13, p=.006), and medication use, REMD, and HF[ms^2^] emerged as significant predictors. Poorer subjective extinction recall was associated with not using medications (B[SE]=-18.81[8.19], p=.024), higher REMD (B[SE]=2.64[.95], p=.007), and lower HF[ms^2^] (B[SE]=-7.05[2.92], p=.018) (**Table 5, Figure S3)**.

**Table 5.**
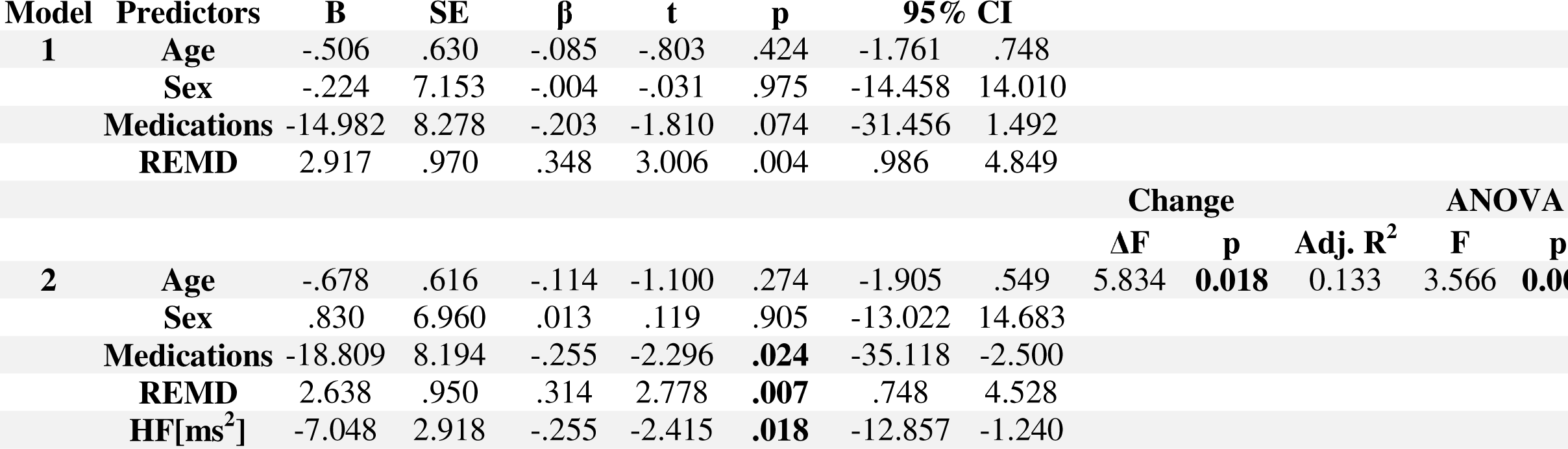
Hierarchical regression analysis for subjective extinction recall (sERI; N=85). Addition of HF[ms^2^] significantly increased the proportion of variance explained by the model. Note that smaller sERI denotes better extinction recall. HF[ms2]: Absolute power of high frequency heart rate variability; REMD: REM density.

For completeness, we repeated the same hierarchical regression analyses by adding all REM variables. The results were similar (**Tables S2 and S3).** Using RMSSD instead of HF[ms^2^] also did not change the results **(Tables S4 and S5).**

## DISCUSSION

We investigated the association of REM measures with the consolidation of physiological and subjective extinction memory in a large sample of trauma-exposed individuals. Confirming our first hypothesis, we found that less time spent in REM sleep, shorter REML, and higher REMD independently predicted poorer physiological extinction recall. Similarly, poorer subjective extinction recall was predicted by higher REMD. Furthermore, our second hypothesis, that higher HF HRV during REM would predict greater extinction recall above and beyond REM measures, was supported for subjective extinction recall and partially supported for physiological extinction recall. To the best of our knowledge, this is the first study to show that features of REM are associated with the consolidation of extinction memory in trauma-exposed individuals. In addition, we show here for the first time that vagal activity during a specific sleep stage contributes to the consolidation of an emotional memory.

REM measures predicting extinction recall is in agreement with previous research in healthy individuals (Davidson and Pace-Schott 2020; Pace-Schott et al. 2015a) and in insomnia disorder (Bottary et al. 2020). In healthy individuals, better extinction recall was associated with the presence of REM during a nap (Spoormaker et al. 2010), higher overnight %REM (Pace-Schott et al. 2014), less fragmented REM, and increased REM theta power (Bottary et al. 2020). In addition, late-night (REM-rich) sleep but not early-night sleep benefited extinction memory (Menz et al. 2016), and selective REM, but not NREM, deprivation impaired extinction recall (Spoormaker et al. 2012). In contrast, among individuals with insomnia disorder (Bottary et al. 2020), better extinction recall was associated with less %REM, shorter REM bouts, and longer REML. Findings are also in agreement with the general notion that REM is involved in processing emotional memories (Goldstein and Walker 2014), although this assertion is not consistently supported by experimental evidence (Davidson et al. 2021).

We further showed that specific REM features were independently associated with extinction recall. There is a sizable literature that implicates both REML and REMD in a variety of psychiatric disorders, including PTSD (Baglioni et al. 2016). However, in PTSD, alterations in these variables are not consistently found, and the direction of change showed variability across studies, possibly due to the heterogeneity in samples and the settings in which data were collected (Baglioni et al. 2016; Kobayashi et al. 2007; Zhang et al. 2019). It was also proposed that two competing processes may be at play simultaneously, REM dysregulation and the resulting pressure to achieve REM (Mellman 1997), which would explain the contradictory findings across studies (e.g., shorter or longer REML, depending on which of these processes predominate). Nonetheless, shortened REML and increased REMD have emerged as characteristics of PTSD in meta-analyses (Baglioni et al. 2016; Kobayashi et al. 2007; Zhang et al. 2019). Our findings advance insight into the significance of these REM alterations and indicate that they are associated with impaired extinction memory, a mechanism that is considered central to the development of PTSD (Bottary et al. 2023).

The mechanisms underlying REM alterations in PTSD and, by extension, how they might be associated with impaired extinction recall are not clear. However, it was proposed that hyperarousal, characterized by impairment in the inhibitory control of amygdala activity by the medial PFC (mPFC) with a concomitant increase in noradrenergic activity, contributes to REM dysregulation (Cabrera et al. 2024; Germain et al. 2008; Pace-Schott et al. 2015b; Pace-Schott et al. 2023). Indeed, increased REMD may be a direct manifestation of hyperarousal in PTSD (Barbato 2023). Rapid eye movements are associated with activation in the limbic and paralimbic structures (Andrillon et al. 2015; Calvo and Fernandez-Guardiola 1984; Corsi-Cabrera et al. 2016; Ioannides et al. 2004; Miyauchi et al. 2009; Wehrle et al. 2007), as well as heart rate surges (Rowe et al. 1999) and nightmares concurrent with autonomic arousal (Paul et al. 2019). Furthermore, in an overlapping sample, we recently showed that REMD was one of the predictors of self-reported hyperarousal symptoms in trauma-exposed individuals (Daffre et al. 2023). Therefore, our results suggest that hyperarousal during REM interferes with the consolidation of extinction memory. REML has been considered an indicator of REM pressure, and the shortening of REML in PTSD has been attributed to an “unmet need” (Mellman 1997). This interpretation would suggest that a history of insufficient REM or a latent factor associated with a REM deficit may impair the consolidation of extinction memory. Alternatively, this association may be due to an adaptive REM enhancement to facilitate emotional processing (Mellman 1997). Yet another possibility is that shortened REML may reflect depression (Palagini et al. 2013). However, self-reported depressive symptoms were not correlated with REML (r_s_=.05, p=.65) or dERI (r_s_=.02, p=.86).

Vagal activity during REM, indexed as HF HRV, was a significant predictor of subjective extinction recall. This novel finding is consistent with the role of the vagal nerve in supporting memory formation (McGaugh 2013) and emotional regulation (Thayer and Lane 2000), as well as the growing body of evidence indicating that vagal activity is causally involved in fear processing (Alvarez-Dieppa et al. 2016; Pena et al. 2014; Pena et al. 2013). Vagal activity in response to cognitive and emotional demands and vagally mediated HRV are considered to reflect the tonic inhibitory control of the amygdala by the mPFC (Thayer et al. 2012; Thayer and Lane 2009), circuitry that is also critical for fear extinction (Milad and Quirk 2012). Consistent with this, high vagal activity during wake was repeatedly shown to be associated with better fear extinction in humans (Jenness et al. 2019; Pappens et al. 2014; Wendt et al. 2019; Wendt et al. 2015). Furthermore, vagal nerve stimulation (VNS) facilitated plasticity in ventromedial PFC-amygdala connectivity in rodents (Pena et al. 2014), and improved extinction learning and recall in both rodents (Alvarez-Dieppa et al. 2016; NobleChuah et al. 2019; Noble et al. 2017; Pena et al. 2014; Pena et al. 2013; SouzaOleksiak et al. 2021; Souza et al. 2022; Souza et al. 2020; SouzaRobertson et al. 2021; Souza et al. 2019) and humans (Burger et al. 2019; Burger et al. 2017; Burger et al. 2016; Szeska et al. 2020). Our results suggest that the contribution of vagal activity continues beyond extinction learning to its consolidation during sleep and supports the potential utility of interventions that can enhance sleep vagal activity in the prevention and treatment of fear-related disorders (NobleSouza et al. 2019). The association of HRV with physiological extinction recall in males should be treated as a preliminary finding, because of the small number of male participants in our sample (N=34). Nevertheless, if replicated, it would be in line with prior research suggesting that sex modulates fear extinction (Milad et al. 2006; Milad et al. 2010; Shvil et al. 2014; Velasco et al. 2019) and its processing during sleep (Schenker et al. 2022).

Our study had several limitations. We had to exclude a substantial number of participants because of lost or unusable sleep data. Lack of oversight during ambulatory recordings can lead to more artifacts than in laboratory studies. Second, there is significant variability across studies in fear conditioning/extinction protocols and methods in analyzing generated data (Lonsdorf et al. 2017). Nonetheless, we used the widely employed Milad et al. 2007 fear conditioning and extinction protocol (Milad and Quirk 2012), and we attempted to be consistent with our own and others’ previously used metrics to index extinction recall. Third, a portion of our sample did not achieve physiological fear conditioning and was therefore excluded from the analysis. Nonetheless, the proportion of “non-conditioners” in our sample (49/139, 35%) was smaller than in many previous studies (Alexandra Kredlow et al. 2018). In addition, we found similar results with the expectancy ratings. A strength of our study is the large sample size. Many of the prior studies linking REM features to emotional processing and extinction were based on small sample sizes, a limitation that reduces statistical power and can lead to Type 1 error and overestimation of effect sizes (Button et al. 2013).

## Conclusions

Abnormalities in REM have repeatedly been reported in individuals diagnosed with PTSD and have been shown to be associated with an increased risk of developing PTSD after a traumatic event. Results of this study further our insight into the role of REM disruptions and indicate that they are associated with impaired consolidation of extinction memory, a mechanism proposed to be critical in the pathogenesis of PTSD.

## Acknowledgements

The authors would like to thank Karen Gannon, RPSGT for scoring sleep recordings and Catherine Bostian for adapting the Yetton et al. REM density program.

## Author Contributions

Edward F. Pace-Schott obtained funding for the study. Lauren Watford, Monami Muranaka, Emma McCoy, Augustus Kram Mendelsohn, Katelyn I. Oliver, Carolina Daffre, Hannah Lax, and Uriel Martinez, collected and reduced data. Cagri Yuksel, Lauren Watford, Monami Muranaka, Hannah Lax, Alexis Acosta, and Abegail Vidrin analyzed the data. Natasha B. Lasko interviewed participants. Scott Orr provided consultation on statistical analyses. Cagri Yuksel and Edward F. Pace-Schott authored the manuscript.

