## Supplementary Materials for "REM disruption and REM Vagal Activity Predict Extinction Recall in Trauma-Exposed Individuals"

**Inclusion and Exclusion Criteria**

All participants reported experiencing a DSM-5 criterion-A traumatic event (“index trauma”) in the past two years, except within the last one month. Participants were allowed with a concurrent anxiety disorder, dysthymia, Major Depressive Disorder (remitted or on case-by-case basis), or if on a stable dose of an antidepressant (≥8 weeks). Exclusion criteria included: history of chronic childhood abuse or neglect; PTSD diagnosis preceding traumatic event indexed at study interview; neurological disorder or injury; major medical disorders; psychotic, bipolar, autism spectrum or other neurodevelopmental disorders; current drug or alcohol abuse or dependence; history of sleep disorder other than insomnia or nightmare disorder; current use of hypnotic or recently adjusted psychiatric medications; shift work; and any contraindication to MRI scans.

### **Ambulatory PSG**

Ambulatory PSG was recorded on 3 nights using the Somte-PSG ambulatory sleep monitor (Compumedics USA, Charlotte, NC, USA). Sampling rate was 256 Hz. EEG data were acquired using six EEG channels (F3, F4, C3, C4, O1, O2; positioned according to the 10-20 system). Additional electrodes were placed on bilateral mastoids, above the right and below the left eye (EOG), under the chin (EMG), and below the right clavicle and in the left fifth intercostal space (ECG). Participants returned home to sleep after being instrumented. During the acclimation/screening (first) PSG night, additional channels for pulse-oximeter, respiration transducer belts, nasal cannula and tibialis movement sensors were added to screen for obstructive sleep apnea (OSA) and Periodic Limb Movement Disorder (PLMD). No participant met criteria for clinically significant OSA or PLMD. All sleep records were scored by an experienced, registered polysomnographic technologist according to American Academy of Sleep Medicine criteria [113].

### **Fear conditioning, extinction learning, and extinction recall procedures**

A well validated 2-day paradigm [60] was used to probe fear conditioning, extinction learning, and extinction memory during ongoing fMRI recording. This protocol consisted of 4 phases, with Habituation, Fear Conditioning, and Extinction Learning phases taking place on the first day and Extinction Recall 24-hours later. During each phase, images of a colored desk lamp (red, yellow, or blue) appearing in a contextual background (office for conditioning context and conference room for extinction context) served as conditioned stimuli (CS). Context images were presented for nine seconds, with three seconds with the lamp off and six seconds with the lamp on (red, yellow or blue). The unconditioned stimulus (US) was a mild (0.8-4.0 mA), 500 msec electric shock delivered to the index and middle fingers of participants’ right hand using a Coulbourn Transcutaneous Aversive Finger Stimulator (Coulbourn Instruments, Allentown, PA). Prior to entering the scanner, participants were administered increasing intensities of shock and they each selected a level that they perceived as “highly annoying but not painful” [64].

During Habituation, all six possible combinations of lamp colors and contexts were presented across six trials. During the following Fear Conditioning phase, two of the three colored lamps (CS+) were each presented 8 times paired with the US at stimulus offset, on a partial reinforcement schedule (5 out of 8 presentations were paired with US). The third lamp color, which was never paired with US (CS-), was interspersed among the CS+s for a total of 16 presentations. Fear Conditioning was followed by Extinction Learning, during which one CS+ (CS+E) was presented in the extinction context 16 times without the US along with 16 interspersed presentations of the CS-. The other CS+ remained conditioned but unextinguished (CS+U). During Extinction Recall, which took place 24 hours later, each CS+ was presented 8 times in the extinction context, with no US, along with 16 interspersed CS-.

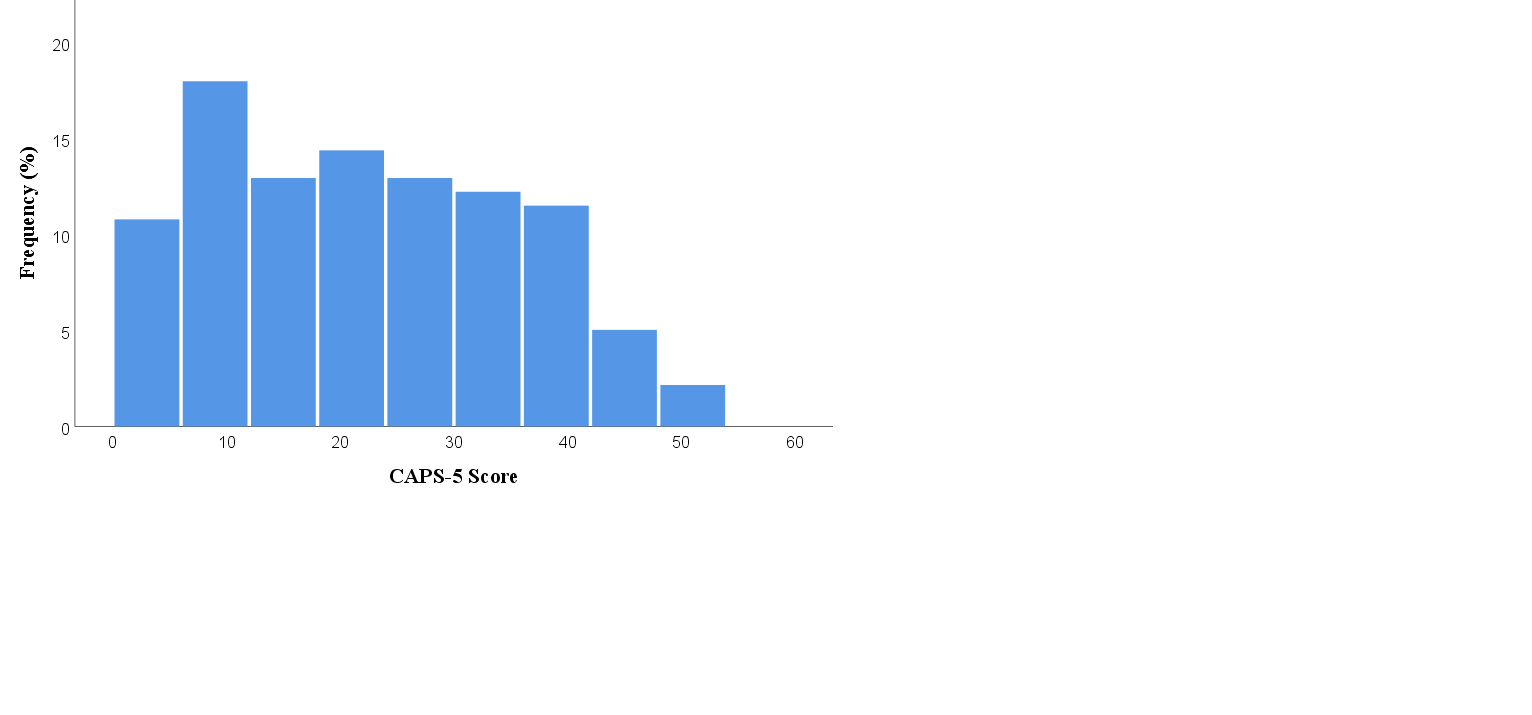

**Figure S1.** Distribution of CAPS-5 score in the sample.

| **Age** | 24.0±4.8 (18-39) |
| --- | --- |
| **Sex (%female)** | 69.9% |
| **Race (%)** |  |
| American Indian  or Alaskan Native | 2.7% |
| Asian | 9.8% |
| Black or African-American | 16.8% |
| More than one race | 6.2% |
| Unknown/unreported | 1.8% |
| White | 62.9% |
| **Ethnicity** |  |
| Hispanic or Latino | 9.7% |
| Not Hispanic or Latino | 85% |
| Unknow/unreported | 3.5% |
| **Type of trauma** |  |
| Transportation accident | 26.5% |
| Violent assault | 19.5% |
| Rape or sexual assault | 17.7% |
| Mass shooting | 2.7% |
| Sudden loss of family or friend | 4.5% |
| Combat incident | 2.7% |
| Multiple or other | 26.4% |
| **Trauma Severity** |  |
| CAPS-5 | 21.8±13.0 (0-53) |
| **Depression Severity** |  |
| QIDS | 7.4±4.4 (0-18) |
| **Months since trauma** | 13.0±6.8 (1-28) |
| **Medications** |  |
| Antidepressants | 18.6% |
| Benzodiazepines | 3.5% |
| Beta-blockers | 0% |
| Mood stabilizers | 0% |
| Antipsychotics | 0% |

**Table S1.** Demographic and clinical characteristics of the participants. Some of the data is displayed as mean ± standard deviation (minimum - maximum).

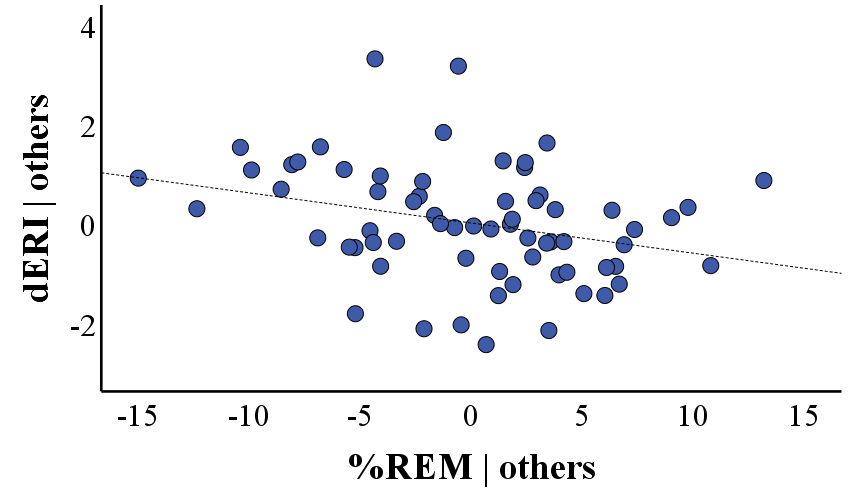

**B**

**A**

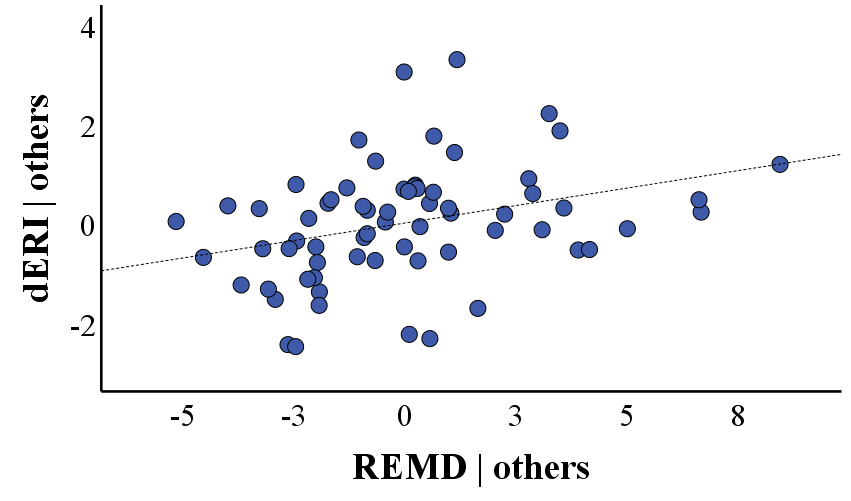

β= .37

β=-.33

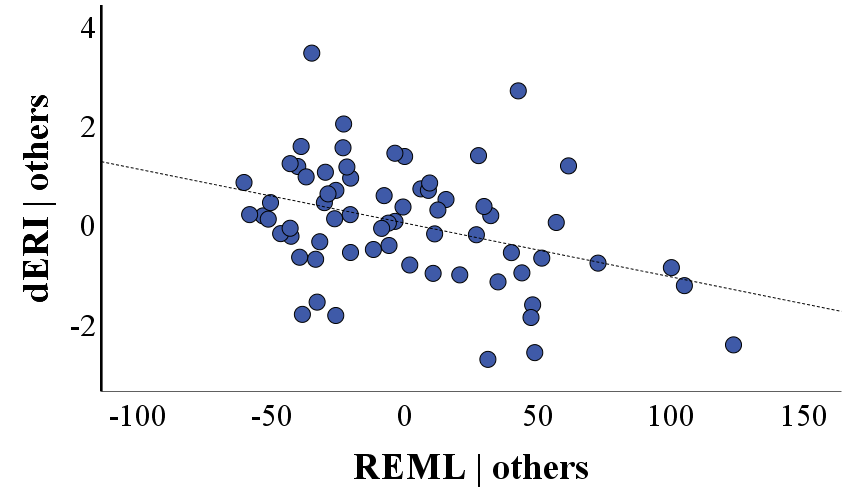

β= -.42

**Figure S2.** Partial regression (added variable) plots of the significant REM variables in the regression analyses that tested the hypothesis 1 for physiological extinction recall (dERI). The Y axis represents the residuals derived from regressing dERI on all the predictor variables in the corresponding model, except the variable noted on the X axis. The X axis represents the residuals derived from regressing the predictor variable noted on the X axis on all the other predictor variables in the corresponding models. The slope reflects the standardized partial regression coefficient (β). Note that smaller dERI denotes better physiological extinction recall. HF[ms^2^]: Absolute power of high frequency heart rate variability; %REM: Proportion of REM sleep to total sleep time; REMD: REM density; REML: REM latency.

**C**

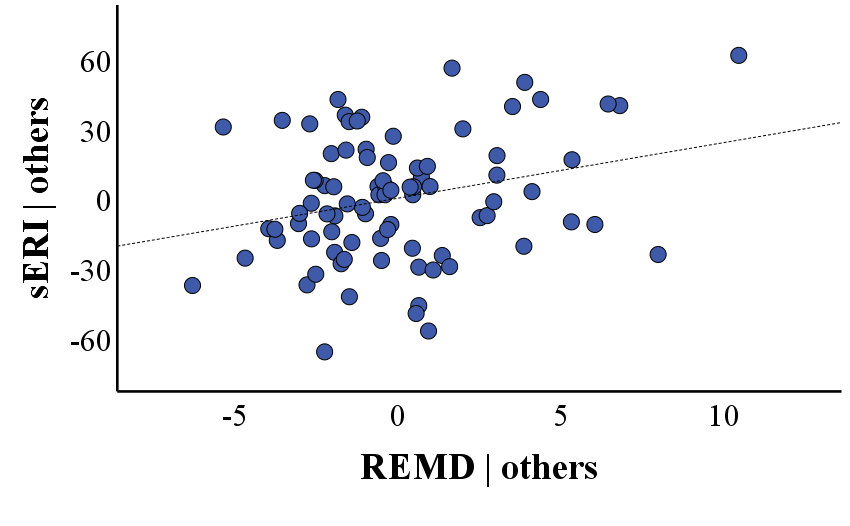

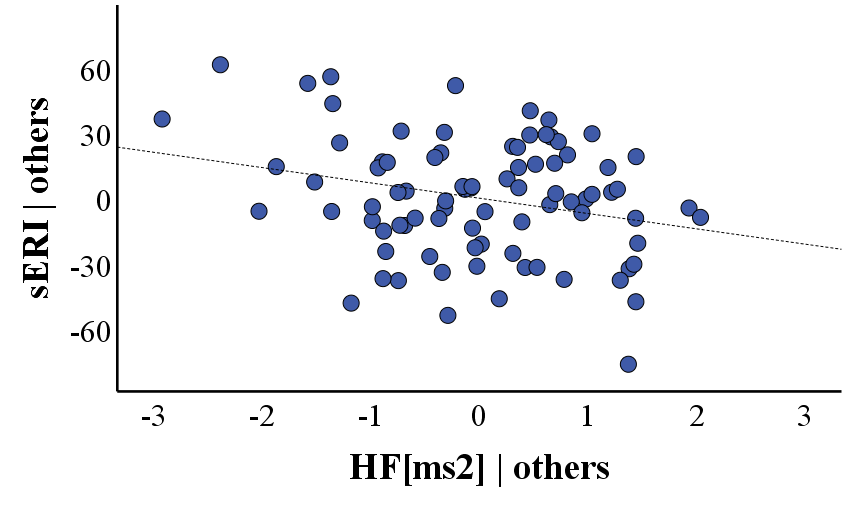

β= -.26

β= .31

**Figure S3.** Partial regression (added variable) plots of the significant REM variables in the regression analyses that tested the hypothesis 1 (A), and the hypothesis 2 (B), for subjective extinction recall (sERI). The Y axis represents the residuals derived from regressing sERI on all the predictor variables in the corresponding model, except the variable noted on the X axis. The X axis represents the residuals derived from regressing the predictor variable noted on the X axis on all the other predictor variables in the corresponding models. The slope reflects the standardized partial regression coefficient (β). Note that smaller sERI denotes better subjective extinction recall. HF[ms^2^]: Absolute power of high frequency heart rate variability; REMD: REM density.

**B**

**A**

| Model | Predictors | B | SE | β | t | p | 95% CI | |  |  |  |  |  |
| --- | --- | --- | --- | --- | --- | --- | --- | --- | --- | --- | --- | --- | --- |
| 1 | **Sex** | -.603 | .351 | -.242 | -1.714 | .092 | -1.307 | .102 |  |  |  |  |  |
|  | **Medication** | -.075 | .396 | -.025 | -.189 | .850 | -.869 | .719 |  |  |  |  |  |
|  | **%REM** | -.052 | .026 | -.281 | -2.004 | .050 | -.104 | .000 |  |  |  |  |  |
|  | **REMD** | .147 | .053 | .391 | 2.793 | .007 | .041 | .253 |  |  |  |  |  |
|  | **REML** | -.010 | .004 | -.383 | -2.688 | .010 | -.017 | -.003 |  |  |  |  |  |
|  | **REMF** | .034 | .032 | .148 | 1.035 | .305 | -.032 | .099 |  |  |  |  |  |
|  |  |  |  |  |  |  |  |  | **Change** | |  | **ANOVA** | |
|  |  |  |  |  |  |  |  |  | **ΔF** | **p** | **Adj. R^2^** | **F** | **p** |
| 2 | **Sex** | -.651 | .345 | -.261 | -1.886 | .065 | -1.344 | .042 | 2.553 | .088 | .162 | 2.450 | **.025** |
|  | **Medication** | .064 | .396 | .022 | .162 | .872 | -.730 | .859 |  |  |  |  |  |
|  | **%REM** | -.058 | .026 | -.313 | -2.280 | **.027** | -.109 | -.007 |  |  |  |  |  |
|  | **REMD** | .159 | .053 | .423 | 3.030 | **.004** | .054 | .265 |  |  |  |  |  |
|  | **REML** | -.012 | .004 | -.446 | -3.122 | **.003** | -.019 | -.004 |  |  |  |  |  |
|  | **REMF** | .028 | .032 | .124 | .885 | .380 | -.036 | .092 |  |  |  |  |  |
|  | **HF[ms^2^]** | -.401 | .201 | -.345 | -1.999 | .051 | -.803 | .002 |  |  |  |  |  |
|  | **HF[ms^2^]×Sex** | .635 | .294 | .391 | 2.160 | .**035** | .045 | 1.224 |  |  |  |  |  |

**Table S2.** Hierarchical regression analysis for physiological extinction recall (dERI) with all REM variables included in the model. Note that smaller dERI denotes better extinction recall. HF[ms^2^]: Absolute power of high frequency heart rate variability; %REM: Proportion of REM sleep to total sleep time; REMD: REM density; REMF: REM fragmentation; REML: REM latency.

| Model | Predictors | B | SE | β | t | p | 95% CI | |  |  |  |  |  |
| --- | --- | --- | --- | --- | --- | --- | --- | --- | --- | --- | --- | --- | --- |
| 1 | **Age** | -.265 | .655 | -.045 | -.404 | .687 | -1.569 | 1.039 |  |  |  |  |  |
|  | **Sex** | .029 | 7.341 | .000 | .004 | .997 | -14.589 | 14.646 |  |  |  |  |  |
|  | **Medication** | -17.151 | 8.405 | -.232 | -2.041 | .045 | -33.886 | -.415 |  |  |  |  |  |
|  | **%REM** | .246 | .543 | .055 | .453 | .652 | -.836 | 1.328 |  |  |  |  |  |
|  | **REMD** | 2.663 | .993 | .317 | 2.681 | .009 | .685 | 4.640 |  |  |  |  |  |
|  | **REML** | .066 | .078 | .103 | .847 | .400 | -.090 | .222 |  |  |  |  |  |
|  | **REMF** | .823 | .671 | .145 | 1.227 | .224 | -.513 | 2.159 |  |  |  |  |  |
|  |  |  |  |  |  |  |  |  | **Change** | |  | **ANOVA** | |
|  |  |  |  |  |  |  |  |  | **ΔF** | **p** | **Adj. R^2^** | **F** | **p** |
| 2 | **Age** | -.433 | .640 | -.073 | -.676 | .501 | -1.707 | .842 |  |  |  |  |  |
|  | **Sex** | 1.142 | 7.143 | .018 | .160 | .873 | -13.083 | 15.368 | 5.681 | **.020** | .137 | 2.671 | **.012** |
|  | **Medication** | -20.916 | 8.312 | -.283 | -2.516 | **.014** | -37.471 | -4.362 |  |  |  |  |  |
|  | **%REM** | .217 | .528 | .048 | .411 | .682 | -.834 | 1.268 |  |  |  |  |  |
|  | **REMD** | 2.399 | .971 | .286 | 2.472 | **.016** | .466 | 4.332 |  |  |  |  |  |
|  | **REML** | .067 | .076 | .103 | .877 | .383 | -.085 | .218 |  |  |  |  |  |
|  | **REMF** | .800 | .651 | .141 | 1.228 | .223 | -.498 | 2.097 |  |  |  |  |  |
|  | **HF[ms^2^]** | -6.940 | 2.912 | -.251 | -2.384 | **.020** | -12.740 | -1.141 |  |  |  |  |  |

**Table S3.** Hierarchical regression analysis for subjective extinction recall (sERI) with all REM variables included in the model. Note that smaller sERI denotes better extinction recall. HF[ms^2^]: Absolute power of high frequency heart rate variability; %REM: Proportion of REM sleep to total sleep time; REMD: REM density; REMF: REM fragmentation; REML: REM latency.

| Model | Predictors | B | SE | β | t | p | 95% CI | |  |  |  |  |  |
| --- | --- | --- | --- | --- | --- | --- | --- | --- | --- | --- | --- | --- | --- |
| 1 | **Sex** | -.531 | .345 | -.213 | -1.539 | .130 | -1.221 | .160 |  |  |  |  |  |
|  | **Medication** | .017 | .386 | .006 | .045 | .964 | -.757 | .791 |  |  |  |  |  |
|  | **%REM** | -.043 | .025 | -.234 | -1.763 | .083 | -.093 | .006 |  |  |  |  |  |
|  | **REMD** | .143 | .053 | .379 | 2.718 | .009 | .037 | .248 |  |  |  |  |  |
|  | **REML** | -.008 | .003 | -.323 | -2.481 | .016 | -.015 | -.002 |  |  |  |  |  |
|  |  |  |  |  |  |  | |  | **Change** | | **ANOVA** | | |
|  |  |  |  |  |  |  |  |  | **ΔF** | **p** | **Adj. R^2^** | **F** | **p** |
| 2 | **Sex** | -.581 | .337 | -.233 | -1.726 | .090 | -1.256 | .094 | 2.700 | 0.076 | 0.164 | 2.686 | **0.019** |
|  | **Medication** | .088 | .382 | .030 | .231 | .818 | -.679 | .855 |  |  |  |  |  |
|  | **%REM** | -.049 | .024 | -.266 | -2.051 | **.045** | -.098 | -.001 |  |  |  |  |  |
|  | **REMD** | .151 | .052 | .402 | 2.914 | **.005** | .047 | .256 |  |  |  |  |  |
|  | **REML** | -.010 | .003 | -.390 | -2.981 | **.004** | -.017 | -.003 |  |  |  |  |  |
|  | **RMSSD** | -.872 | .396 | -.375 | -2.200 | **.032** | -1.667 | -.077 |  |  |  |  |  |
|  | **RMSSD×Sex** | 1.209 | .580 | .369 | 2.085 | **.042** | .046 | 2.371 |  |  |  |  |  |

**Table S4.** Hierarchical regression analysis for physiological extinction recall (dERI). RMSSD and RMSSD × Sex interaction were significant predictors. Note that smaller dERI denotes better extinction recall. %REM: Proportion of REM sleep to total sleep time; REMD: REM density; REML: REM latency; RMSSD: Root mean square of successive differences.

| Model | Predictors | B | SE | β | t | p | 95% CI | |  |  |  |  |  |
| --- | --- | --- | --- | --- | --- | --- | --- | --- | --- | --- | --- | --- | --- |
| 1 | **Age** | -.506 | .630 | -.085 | -.803 | .424 | -1.761 | .748 |  |  |  |  |  |
|  | **Sex** | -.224 | 7.153 | -.004 | -.031 | .975 | -14.458 | 14.010 |  |  |  |  |  |
|  | **Medication** | -14.982 | 8.278 | -.203 | -1.810 | .074 | -31.456 | 1.492 |  |  |  |  |  |
|  | **REMD** | 2.917 | .970 | .348 | 3.006 | .004 | .986 | 4.849 |  |  |  |  |  |
|  |  |  |  |  |  |  | |  | **Change** | | **ANOVA** | | |
|  |  |  |  |  |  |  |  |  | **ΔF** | **p** | **Adj. R^2^** | **F** | **p** |
| 2 | **Age** | -.628 | .620 | -.106 | -1.013 | .314 | -1.861 | .605 | 4.515 | **0.037** | 0.133 | 3.265 | **0.010** |
|  | **Sex** | .062 | 7.002 | .001 | .009 | .993 | -13.875 | 13.999 |  |  |  |  |  |
|  | **Medication** | -18.339 | 8.255 | -.248 | -2.222 | **.029** | -34.770 | -1.909 |  |  |  |  |  |
|  | **REMD** | 2.758 | .953 | .329 | 2.895 | **.005** | .862 | 4.655 |  |  |  |  |  |
|  | **RMSSD** | -12.436 | 5.853 | -.225 | -2.125 | **.037** | -24.085 | -.787 |  |  |  |  |  |

**Table S5.** Hierarchical regression analysis for subjective extinction recall (sERI). Addition of RMSSD significantly increased the proportion of variance explained by the model. Note that smaller sERI denotes better extinction recall. REMD: REM density; RMSSD: Root mean square of successive differences.
